# Discovery and engineering of AiEvo2, a novel Cas12a nuclease for human gene editing applications

**DOI:** 10.1101/2023.08.28.554880

**Authors:** Allison Sharrar, Luisa Arake de Tacca, Zuriah Meacham, Johanna Staples-Ager, Trevor Collingwood, David Rabuka, Michael Schelle

## Abstract

The precision of gene editing technology is critical to creating safe and effective therapies for treating human disease. While the programmability of CRISPR-Cas systems has allowed for rapid innovation of new gene editing techniques, the off-target activity of these enzymes has hampered clinical development for novel therapeutics. Here we report the identification and characterization of a novel CRISPR-Cas12a enzyme from *Acinetobacter indicus* (AiCas12a). We then engineer the nuclease (termed AiEvo2) for increased specificity, PAM recognition, and efficacy on a variety of human clinical targets. AiEvo2 is highly precise and able to efficiently discriminate between normal and disease-causing alleles in Huntington’s patient derived cells by taking advantage of a single nucleotide polymorphism on the disease-associated allele. AiEvo2 efficiently edits several liver-associated target genes including PCSK9 and TTR when delivered to primary hepatocytes as mRNA encapsulated in a lipid nanoparticle. The enzyme also engineers an effective CD19 CAR-T therapy from primary human T cells using multiplexed simultaneous editing and CAR insertion. To further ensure precise editing, we engineered an anti-CRISPR protein (ErAcr) to selectively inhibit off-target gene editing while retaining therapeutic on-target editing. The engineered AiEvo2 nuclease coupled with a novel ErAcr protein represents a new way to control the fidelity of editing and improve the safety and efficacy of gene editing therapies.

## Introduction

Since the discovery and development of Cas9 as a tool for human gene editing therapies, multiple CRISPR nucleases have emerged offering alternative optionality for drug developers. The type V-A CRISPR family (Cas12a) has emerged as the clear alternative to Cas9 for therapeutic applications. The Cas12a nucleases offer several advantages for human gene editing, including a shorter guide RNA (∼40 nucleotides vs >100 for Cas9), the ability to process a guide array into multiple targeting guides, and reduced off-target editing due to the intrinsic precision of the enzyme^1,2^.

Off-target gene editing is a key concern for any CRISPR therapy. While increased editing precision is desirable for all gene editing therapies, the need is particularly acute with *in vivo* interventions that rely on delivery of the editing technology directly to affected cells in the patient, eliminating the ability to screen for unintended adverse editing outcomes. Recently, the US Food and Drug Administration weighed in for the first time on the risks of off-target editing and outlined a series of tests required for any new CRISPR therapy to advance toward commercialization^3^. As the therapeutic gene editing field advances, acceptable off-target risks will continue to decrease, putting pressure on the field to increase editing precision.

We set out to discover novel Cas12a enzymes and engineer them for use in human therapeutics. This led to the discovery of 98 new Cas12a effector candidates from diverse bacterial and metagenomic sources. These candidates were screened for gene editing in human cells. From this library, we characterize and engineer AiCas12a from *Acinetobacter indicus*. The engineered enzyme, termed AiEvo2, has broad PAM recognition, enhanced activity, and high specificity. We demonstrate the efficacy of AiEvo2 for therapeutic applications with allele specific editing for Huntington’s disease, editing of liver-associated targets PCSK9 and TTR, and generation of a CD19 CAR-T from primary human T cells. Finally, we increase the overall precision of CRISPR gene editing by identification and engineering of a novel anti-CRISPR protein active against AiCas12a. The engineered anti-CRISPR (ErAcr) selectively reduces off-target editing while retaining therapeutic on-target efficacy. Collectively, this work highlights a new CRISPR gene editing system comprising a novel, engineered AiEvo2 nuclease with broad PAM recognition and an engineered ErAcr protein that increases editing precision for therapeutic applications.

## Results

### Discovery of 98 unique Cas12a candidates

We mined several metagenomic and protein databases for novel Cas12a sequences using hidden Markov models consisting of known Cas12a family members. To bias our candidates toward sequences that would be functional in human cells, we selected candidates that contained complete RuvC domains and key active site residues. We further pared the candidate list by only including sequences from genomes with an associated CRISPR array and identifiable direct repeat region. This resulted in the identification of 98 unique CRISPR-Cas12a candidates with identities <90% to known Cas12a enzymes^4–7^. Each Cas12a candidate was paired with its cognate direct repeat sequence. The 98 Cas12a candidates use a combined 16 unique direct repeat sequences ranging from 19-22 nucleotide in length.

### Identification of 67 novel Cas12a nucleases active in human cells

The 98 Cas12a candidates were codon-optimized for human cell expression and cloned into mammalian expression vectors along with a single guide RNA (sgRNA) consisting of the cognate direct repeat and a spacer for a human gene target. The candidates were then tested for gene editing in human cells. Over two thirds (67 of 98) of the candidates had detectable gene editing (>0.1%) in human cells (Figure 1A), highlighting the efficiency of our bioinformatics approach and the requirement for a cognate direct repeat sequence. We focused our initial development around AiCas12a, a unique Cas12a candidate identified in the soil bacterium *Acinetobacter indicus* isolated from a hexachlorocyclohexane dump site^8^. AiCas12a is distinct from the canonical Cas12a enzymes AsCas12a and LbCas12a ^9^, sharing <40% identity to either protein. Also, AiCas12a has an extra nucleotide in its cognate direct repeat when compared to AsCas12a or FnCas12a (20 versus 19 nucleotides), resulting in an expanded loop similar to the CRISPR RNA (crRNA) of MbCas12a and LbCas12a (Figure S1).

**Figure 1.**
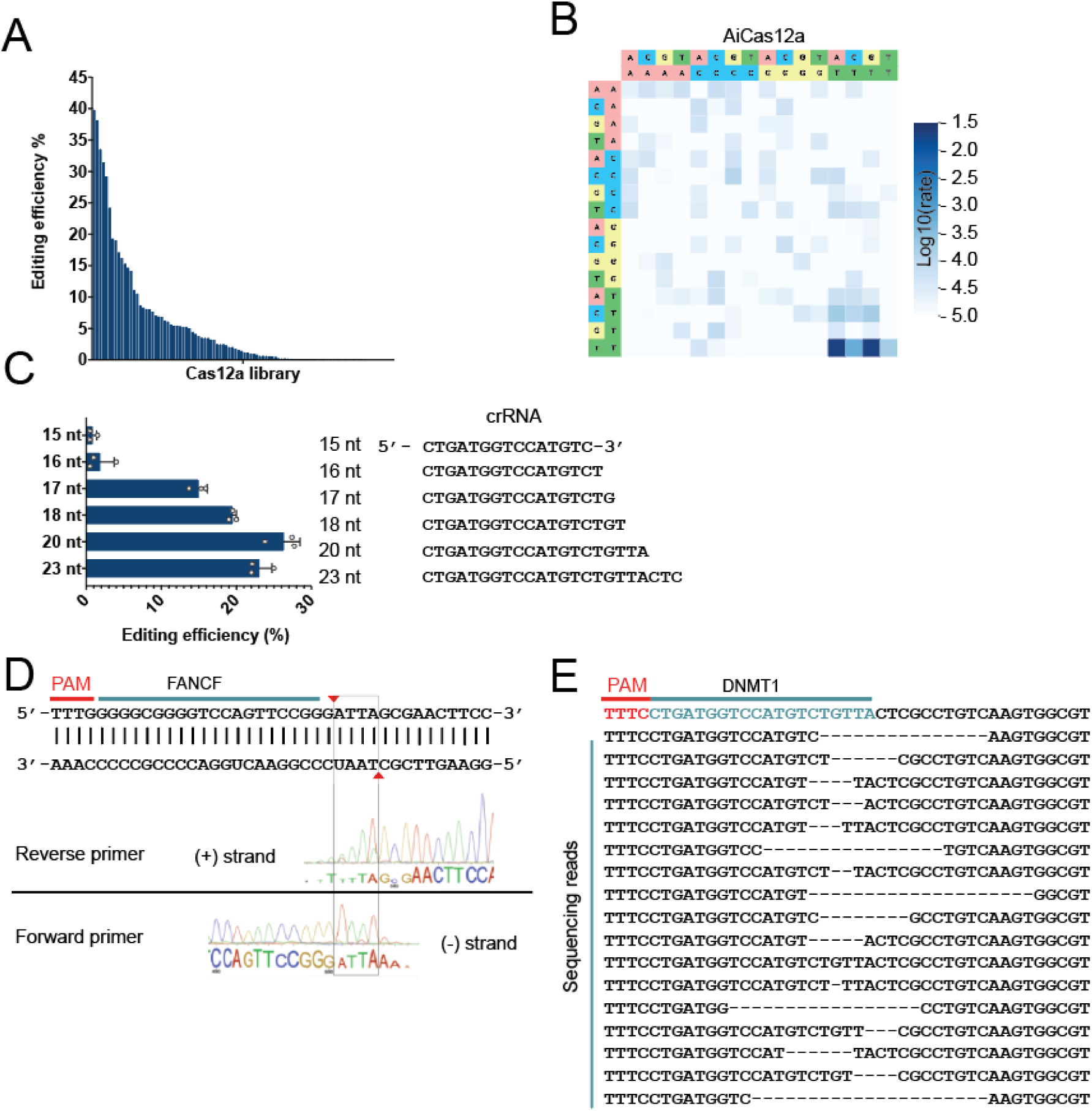
Identification and Characterization of AiCas12a. **(A)** Editing efficiencies of 98 novel Cas12a nucleases in HEK293T human cells. 67 of the 98-member library showed detectable editing by plasmid transfection, including AiCas12a. **(B)** PAM characterization of AiCas12a showing preference for TTTV PAM sequences. **(C)** Spacer length characterization for AiCas12a showing a preference for a 20-nucleotide spacer. Editing was performed using plasmid transfection of HEK293T cells and analyzed by Sanger sequencing and TIDE analysis. N = 3 biological replicates. **(D)** Cleavage analysis of AiCas12a. Cleavage sites on FANCF are indicated in red arrows. Sanger traces show cleavage 19 nucleotides from the PAM on non-target strand and 23 nucleotides from the PAM on the target strand. The (–) strand sequence is reverse- complemented to show the top strand sequence. **(E)** Next generation sequencing reads showing representative indel pattern for AiCas12a targeting DNMT1. Deletions range from nucleotide 7 to nucleotide 34 distal to the PAM.

### Native AiCas12a prefers thymine-rich PAM sequences and a 20-nucleotide spacer

The protospacer adjacent motif (PAM) is an important characteristic of any CRISPR-Cas nuclease. By recognizing the PAM, the CRISPR-Cas system is able to make a distinction between self and non-self and cleave the target DNA. In therapeutic applications, the PAM often restricts editing to particular genomic locations that have the preferred PAM for the given nuclease, while other locations remain unreachable. Cas12a nucleases most commonly prefer a thymine-rich, four base PAM (TTTV) that is found adjacent to the 5′-end of the spacer. We employed the HT-PAMDA assay^11^ to determine the PAM preferences of AiCas12a (Figure 1B). Similar to other members of the Cas12a family, AiCas12a prefers a 5′ TTTV PAM and shows maximal activity with A or G in the fourth position.

Spacer length is another key component in optimizing CRISPR-Cas systems for human gene editing applications. SpyCas9 prefers a 20-23 nucleotide spacer ^12^ while SauCas9 prefers a 21- nucleotide spacer^13^. Previously reported Cas12a nucleases were found to prefer a 20-23 nucleotide spacer^2,4^. We tested AiCas12a with spacers ranging from 15-23 nucleotides in length (Figure 1C). Similar to other Cas12a nucleases, AiCas12a preferred spacers between 18-23 nucleotides and optimal editing efficiency with a 20-nucleotide spacer. Further truncation of the spacer (15-16 nucleotides) resulted in a dramatic loss of activity.

### AiCas12a cleaves dsDNA leaving a staggered 5′ overhang

We next mapped the cleavage site of AiCas12a using *in vitro* cleavage of linear dsDNA followed by Sanger sequencing on the DNA ends (Figure 1D). AiCas12a cleavage leads to a 4-nucleotide 5′ overhang similar to the cleavage pattern for other Cas12a family nucleases^4^. The cut site is distal to the PAM and occurs after the 19^th^ nucleotide on the (+) strand and after the 23^th^ nucleotide on the (-) strand. We then assessed the editing outcomes of AiCas12a targeting DNMT1 in HEK293T cells using NGS amplicon sequencing (Figure 1E). The indel pattern shows gene editing centered around the 3′ end of the guide, consistent with the cleavage sites identified above. The most common editing outcome is deletion ranging in size from position 7 proximal to PAM to position 34 proximal do PAM, similar to the indel pattern observed for other Cas12a family members ^14^.

### NLS optimization increases editing efficiency in human cells

Effective gene editing in eukaryotic systems requires localizing the Cas enzyme to the nucleus of the cells. Large proteins like CRISPR-Cas nucleases can be fused to nuclear localization signals (NLS) to allow them to reach the nucleus and edit the genomic DNA. Previous studies have shown that the optimal NLS for different families of nucleases is not universal ^15^ . For instance, SpyCas9 with three NLS (1 N-terminus and 2 C-terminus) had increased activity in stem cells. However, AsCas12a activity was maximized with a different combination of signals with just two C-terminal NLS and no N-terminal NLS. When the SpyCas9 configuration of was added to AsCas12a, Cas12a activity was decreased^16^. To identify the optimal NLS configuration for AiCas12a, we assessed six different NLS configurations consisting of four distinct NLS sequences (SV40, c- Myc, nucleoplasmin, and a longer SV40 peptide; Figure 2A). We observed the highest editing using the longer SV40 peptide NLS on the C-terminus of the nuclease (Figure 2C). Similar to the study cited above, adding additional NLS to the N-terminus did not improve editing of AiCas12a (Figure 2B).

**Figure 2.**
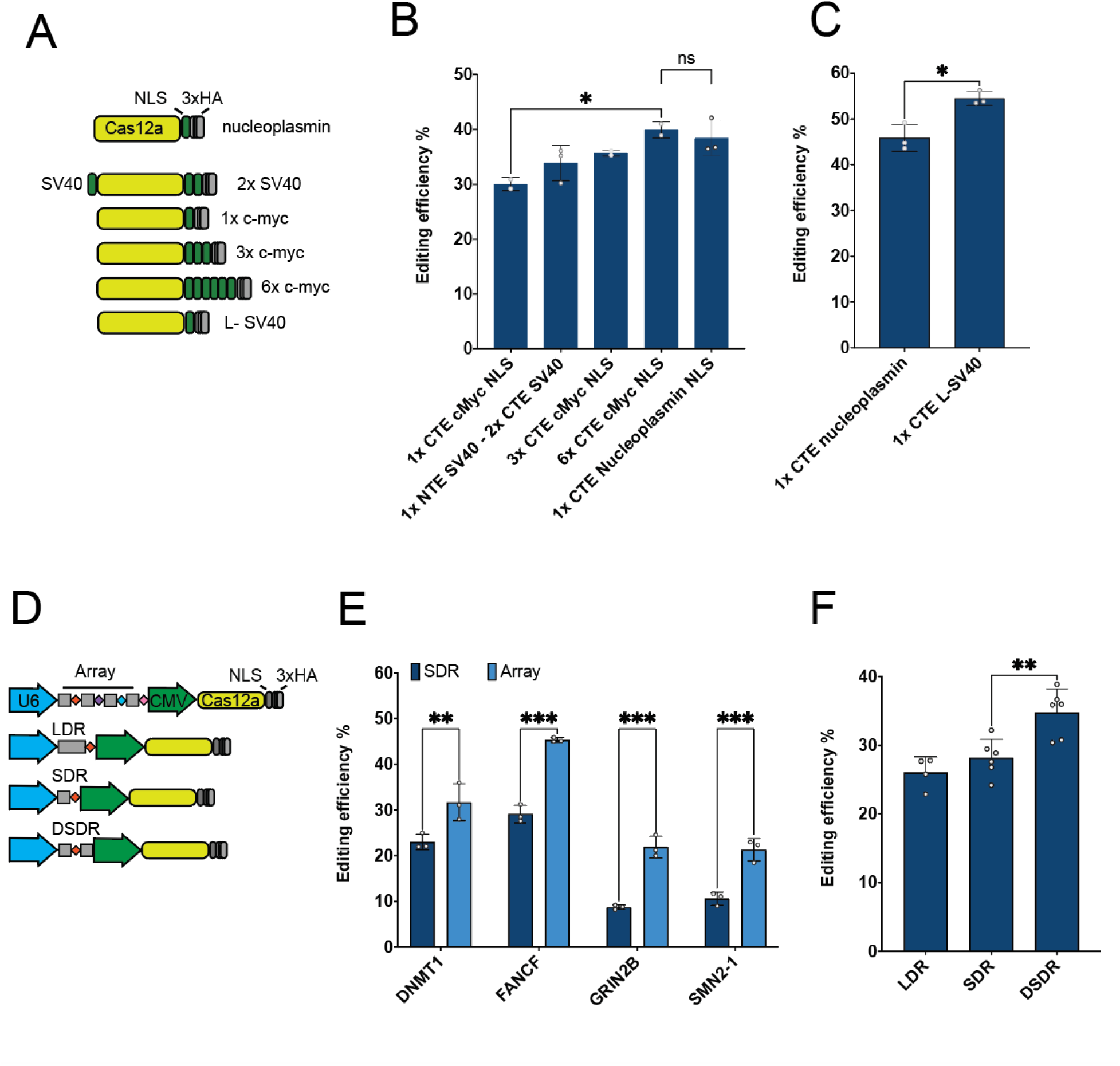
Optimization of AiCas12a for human gene editing. **(A)** Schematic of different Nuclear Localization Signals (NLS) tested with AiCas12a in HEK293T cells. **(B)** Editing of FANCF using NLS configurations shown in panel A. 6x CTE c-Myc and 1x CTE Nucleoplasmin show significantly higher editing than 1x CTE c-Myc (p = 0.04). **(C)** Editing of FANCF using nucleoplasmin and a longer SV40 (L-SV40) NLS configurations. The L-SV40 NLS was significantly more efficient than nucleoplasmin (p = 0.01). **(D)** Schematic of AiCas12a guide RNA configurations. Long direct repeat (LDR), short direct repeat (SDR), double short direct repeat surrounding the spacer (DSDR), guide array with four spacers and direct repeats (Array). **(E)** Editing profiles of four target genes using either a single spacer and short direct repeat (SDR) or an array with all four spacers and four direct repeats (Array). Editing at each site was significantly increased using the Array relative to the SDR configuration (p < 0.007). **(F)** Editing of FANCF with various guide RNA configurations from panel D. The DSDR configuration was significantly higher than either the LDR or SDR configurations. Having two short direct repeat sequences (DSDR) around the spacer is significantly more efficient than having only one direct repeat (DR) sequence upstream of the spacer (p = 0.004). Editing in B, C, E, and F was performed using plasmid transfection of HEK293T cells and analyzed by Sanger sequencing and TIDE analysis. N = 3-6 biological replicates.

### AiCas12a simultaneously edits multiple genes using a multiplexed array

One of the main advantages of Cas12a nucleases is their ability to process their own sgRNA from an array, allowing for multiplexed editing of multiple gene targets ^17,18^. Several different spacers can be added in an array separated by the nuclease-specific direct repeat with expression driven by a single U6 promoter. The nuclease processes each direct repeat and spacer combination into separate guides, allowing simultaneous editing at different genomic locations. To test AiCas12a for multiplexed editing, we constructed an array targeting four genes (DNMT1, FANCF, GRIN2B, and SMN2) and compared the editing of the array versus the individual guides (Figure 2D & E). Editing using the array was significantly higher for each target as compared to editing with the single guide.

We extended these results by testing different guide configurations including the mature crRNA (Short Direct Repeat, SDR), the pre-processed crRNA (Long Direct Repeat, LDR) or a Double Short Direct Repeat (DSDR) with the direct repeat on either side of the spacer (Figure 2D & F). While there was no significant difference between the long and short direct repeat, there was increased activity with the short direct repeat on either side of the spacer. This result combined with the increased editing seen with the array indicates that AiCas12a is more active after processing the guide. This is consistent with previous multiplexing experiments^19^.

### Mismatch editing profile of AiCas12a

The Cas12a family has been shown to have lower off-target editing than Cas9 ^2,20^. To test the potential for off-target activity with native AiCas12a, we designed a mismatch panel by substituting each position in a spacer targeting DNMT1 to its Watson-Crick binding counterpart (Figure 3A). AiCas12a shows very low editing when positions 2 through 7 are mismatched, which overlaps with the predicted seed region ^21^ as well as positions 10-17. At the PAM-distal region of the guide, AiCas12a is less able to discriminate between mismatches of the guide and target sequence, similar to previous results with other Cas12a family members ^2^.

**Figure 3.**
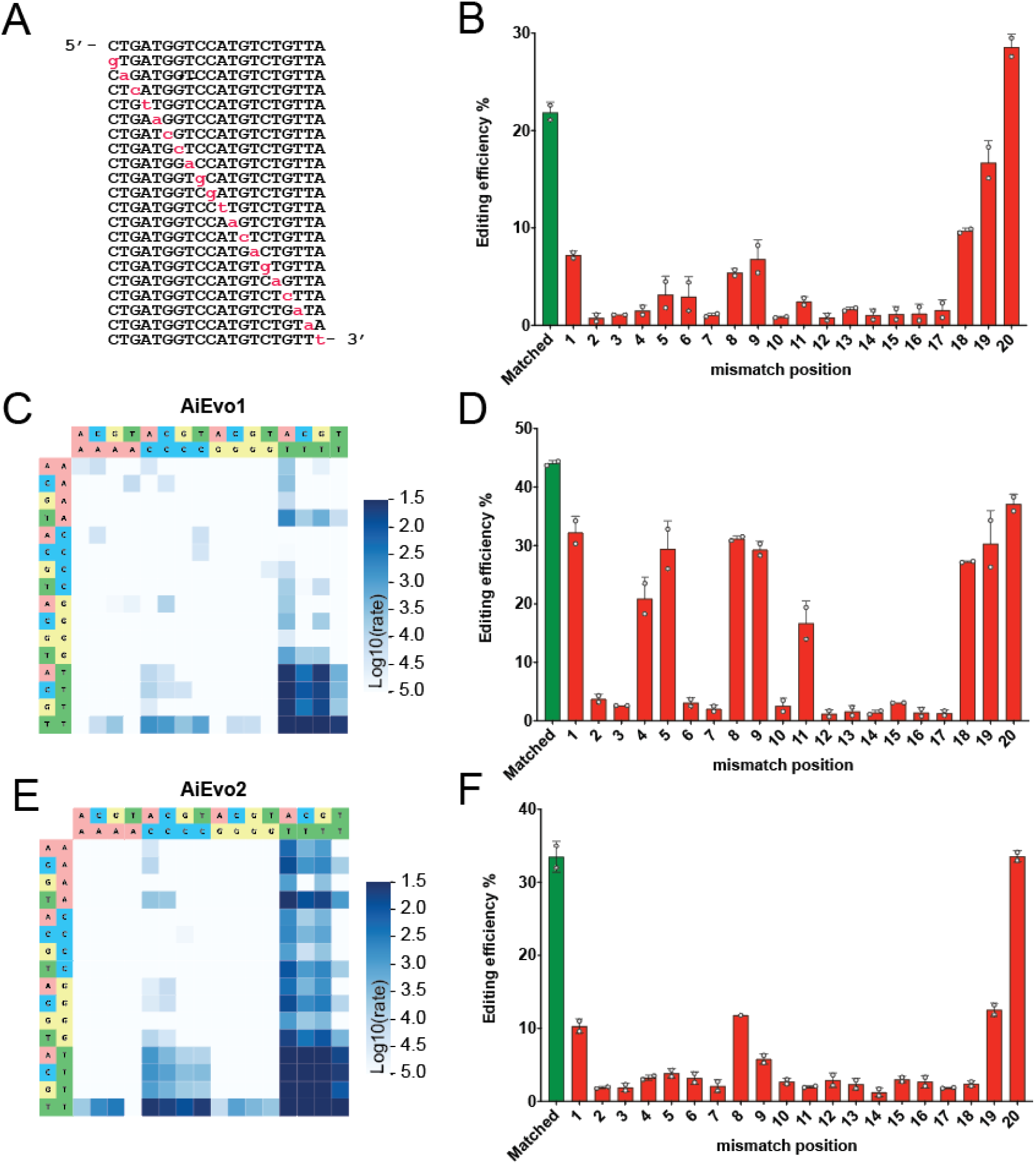
Engineering AiCas12a for broader PAM recognition and fidelity. (**A)** Representation of the DNMT1 single mismatch guide panel. Mutated nucleotides are shown in red. (**B)** Editing profile of AiCas12a with the DNMT1 mismatch panel (red) versus the fully matched guide (green). **(C)** PAM preferences of AiEvo1 showing an expanded PAM relative to native AiCas12a. **(D)** DNMT1 mismatch panel editing profile of AiEvo1. PAM expansion resulted in a loss of specificity with increased editing at several positions. **(E)** PAM preference of AiEvo2 showing a further broadening of PAM preference. **(F)** DNMT1 mismatch panel editing profile of AiEvo2. Further engineered increased the fidelity of AiEvo2 while maintaining PAM diversity. Editing for B, D, and F were performed using plasmid transfection of HEK293T cells and analyzed by Sanger sequencing and TIDE analysis. N = 3 biological replicates.

### AiEvo2, engineered AiCas12a for increased activity, PAM diversity, and fidelity

The ability to target a broad set of genomic targets is required for a robust and widely applicable CRISPR-Cas nuclease. Many therapeutic targets require editing in a small window, making guide design challenging. A limiting factor in guide design is PAM availability. The native TTTV PAM^4,22^ of AiCas12a is fairly restrictive, with a theoretical frequency of 1 PAM site every 42 bases, compared to the 1 PAM site every 8 bases for the NGG PAM of SpyCas9. To increase the PAM diversity of our AiCas12a nuclease, we mutated amino acids around the predicted PAM-interacting domain. This technique has been used to increase the PAM recognition of related enzymes like AsCas12a ^23–25^. Substituting E174R, S598R, and K604R increased the activity of AiCas12a (Figure S2) and dramatically increased the PAM recognition of the engineered enzyme, termed AiEvo1 (Figure 3C).

Previous studies found that increasing the PAM recognition of Cas12a enzymes also increased the promiscuity of editing at mismatch locations^26^. We tested AiEvo1 against the mismatch panel in Figure 3A and found an increase in editing at several mismatch locations, with mismatch 1, 5, 8, and 9 editing at approximately 75% of the fully matched guide (Figure 3D)^11,25,53^. To counteract this loss in precision, we substituted N283A to produce AiEvo2. AiEvo2 maintained the expanded PAM from AiEvo1 and regained the specificity seen in the wild-type enzyme (Figure 3E & F). AiEvo2 efficiently edits a much broader PAM (NTTN) as well as additional PAM sequences like TTCN and TATV. This increases the theoretical genome targeting density from 1 in 42 bases to 1 in every 6 bases, yielding a theoretical density greater than SpyCas9.

### Allele specific editing to treat Huntington’s disease

Huntington’s disease (HD) is characterized by expanded CAG repeats in the huntingtin (*HTT*) gene that lead to mutant protein production. The disease-causing *HTT* gene is typically inherited from a single affected parent and present as a single allele^27–29^. Several single nucleotide variants (SNV) have been associated with the mutant *HTT* allele and have been used as genetic handles to differentiate the mutant and normal allele. Previous attempts to develop CRISPR strategies using *HTT* SNVs relied on variations that altered the PAM, which limits the options for guide design^30^. Given the specificity of AiEvo2, we reasoned that the enzyme would be able discriminate between the mutant and normal allele by using the SNV to create a mismatch within guide sequence. This gives us more guide design options to optimally position the SNV in the guide.

We designed two guides targeting the rs363099 SNV in exon 28 of the *HTT* gene (Figure 4A). The SNV was placed at position 2 for guide GSp340 and position 11 for guide GSp341. As demonstrated in Figure 3F, these guide positions are relatively intolerant of mismatches. We then nucleofected HD patient fibroblasts using ribonucleoprotein (RNP) comprising AiEvo2 and GSp340 or GSp341 (Figure 4B). Both guides are able to edit the mutant allele at high levels (∼80%). While GSp341 shows some editing at the normal allele (∼25%), GSp340 shows near complete selectivity with <1% editing at the normal allele. This indicates that mismatches at guide position 2 offer more selectivity than at position 11 and that targeting a SNV can generate allele specific editing outcomes.

**Figure 4.**
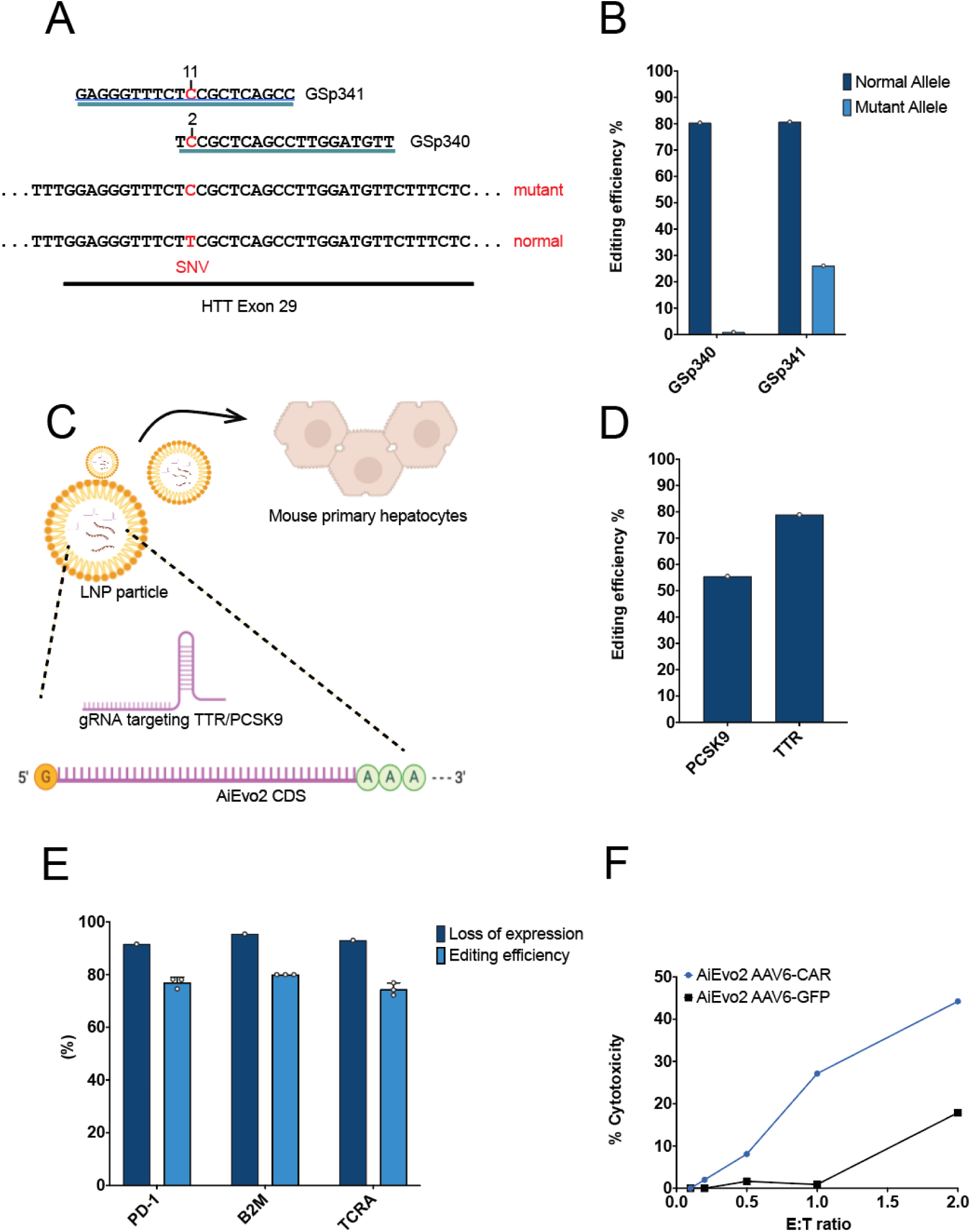
Therapeutic Applications of AiEvo2. **(A)** Graphical representation of the strategy used to selectively edit HD patient alleles containing a SNV. Two guides target the rs363099 SNV in *HTT* exon 28. The SNV creates a mismatch with the normal allele at guide position 2 for GSp340 and at guide position 11 for GSp341. **(B)** Allele specific editing using AiEvo2. Guide GSp340 only edits the mutant allele with <1% editing at the normal allele. **(C)** Graphical representation of LNP-based editing of mouse primary hepatocytes using LNP containing AiEvo2 and guides targeting TTR or PCSK9 genes. **(D)** Editing efficiencies of AiEvo2 at the PCSK9 or TTR genomic locations in mouse primary hepatocytes using lipid nanoparticle delivery. **(E)** Multiplexed triple target editing for CAR-T engineering. Flow cytometry (dark blue) shows >90% decrease of cell surface markers for all three targets while editing efficiencies of show >80% editing at each loci (light blue). Editing was performed by nucleofecting AiCas12a RNP into primary human T cells and analyzed by Sanger sequencing and TIDE analysis. N = 3 biological replicates. **(F)** Demonstration of CAR-T activity. CD19^+^ NALM-6 human cancer cells were treated with increasing CAR-T effector to target ratios (E:T ratio). Evo2Car-T shows increased cytotoxicity relative to control engineered T cells.

### AiEvo2 mRNA delivered by LNP efficiently edits liver disease targets in primary hepatocytes

Lipid nanoparticles (LNP) are a key delivery modality for *in vivo* CRISPR gene editing therapies^31,32^. Typically, mRNA encoding the nuclease is packaged along with sgRNA into LNP, which are effectively up taken by liver hepatocytes. To test our engineered AiEvo2 nuclease for *in vivo* therapies, we developed guides targeting TTR and PCKS9, two validated liver gene targets^33–35^. We produced and packaged mRNA encoding AiEvo2 along with sgRNA targeting either TTR or PCKS9 into LNP (Figure 4C). The LNP were then used to transfect primary mouse hepatocytes. We observed high editing efficiency for both TTR and PCSK9 in primary hepatocytes, indicating that AiEvo2 mRNA is efficiently packaged and delivered to primary cells (Figure 4D).

### Engineering a CD19 CAR-T with AiEvo2

Allogeneic Chimeric Antigen Receptor (CAR) T cell therapies are prime targets for CRISPR-based gene editing techniques^36–38^. Allogeneic CAR-T therapies rely on disruption of multiple genes to reduce the risk of graft-versus-host-disease (GvHD) and increase the potency of the therapeutic. We sought to evaluate AiEvo2 as a tool to generate an allogeneic CAR-T therapy targeting CD19+ cancer cells. We targeted simultaneous disruption of three common T cell targets: PD-1, B2M, and TCRA while inserting a CD19 CAR into the TCRA locus (Figure S3). PD-1 is a critical regulator of the fate and function of T cells. Rupp and collaborators ^39^ showed that PD-L1 in tumor cells was detrimental to CAR-T mediated killing *in vitro* as well as for tumor clearance *in vivo*. Disruption of PD-1 in the T cells increases the potency of the CAR-T therapy. Meanwhile, disruption of TCRA and B2M help alleviate GvHD and increase the longevity of the CAR-T therapy. Targeting the endogenous T cell receptor is a common strategy to eliminate cross reactivity with the allogeneic cell^40^. Beta-2-microglobulin (B2M) is an essential subunit of human leukocyte antigen class I (HLA-I) proteins. The presence of foreign HLA-I on the allogeneic cells leads to rapid elimination and decreased potency. Pre-clinical data shows that the combination of B2M disruption and the knockout of TCRA produces universal CD19-targeting T cells^41^.

To generate a CAR-T therapy with AiEvo2, we first designed a CAR cassette targeting CD19 (Figure S3B)^42^. The CAR was designed with homology arms to the TCRA target site, putting the CAR under endogenous TCRA expression (Figure S4). The CAR cassette was packaged into an AAV6 viral vector and used to transduce primary human T cells. Guides targeting TCRA, B2M, and PD-1 were designed and simultaneously nucleofected along with AiEvo2 protein as RNP into primary human T cells. All three guides show high levels of editing and efficient elimination of the corresponding cell surface protein as evidenced by flow cytometry (Figure 4E and S5).

We then tested the ability of our engineered CAR-T cells to recognize CD19-expressing NALM6 cancer cells. NALM6 cells are a B-cell precursor leukemia cell line which has been shown to express high levels of CD19^43^. We incubated NALM6 cells with our engineered CAR-T cells at various effector to target (E:T) ratios and assessed CAR-T mediated NALM6 cell killing. Our triple knockout CD19 CAR-T cells were able to efficiently eliminate NALM6 cells at a 1:1 E:T ratio relative to control engineered T cells (Figure 4F).

### Engineered anti-CRISPR proteins increase gene editing precision

Having demonstrated AiEvo2 as a robust and broadly applicable gene editing tool, we next focused on ensuring the overall precision of gene editing. We have developed a complementary approach to control gene editing outcomes by engineering anti-CRISPR (Acr) proteins to selectively inhibit off-target editing. Acr proteins evolved in bacteriophage as a way to inhibit bacterial CRISPR defenses^44–46^. Acr proteins have been identified that inhibit many Type II and Type V CRISPR systems, including Cas9 and Cas12a^46–53^. However, previously identified Cas12a anti-CRISPR proteins AcrVA1 and AcrVA5 showed no activity against AiEvo2 in human cells (Figure 5A). To identify Acr proteins that inhibit AiEvo2, we mined metagenomic databases for distant homologs of Cas12a anti-CRISPR AcrVA1 (Figure S6). We identified five candidate Acr proteins (termed ACX175-179) and tested them for inhibition of AiEvo2 in HEK293T cells. Two AcrVA1 homologs, ACX175 and ACX177 fully inhibited AiEvo2 (Figure 5A).

**Figure 5.**
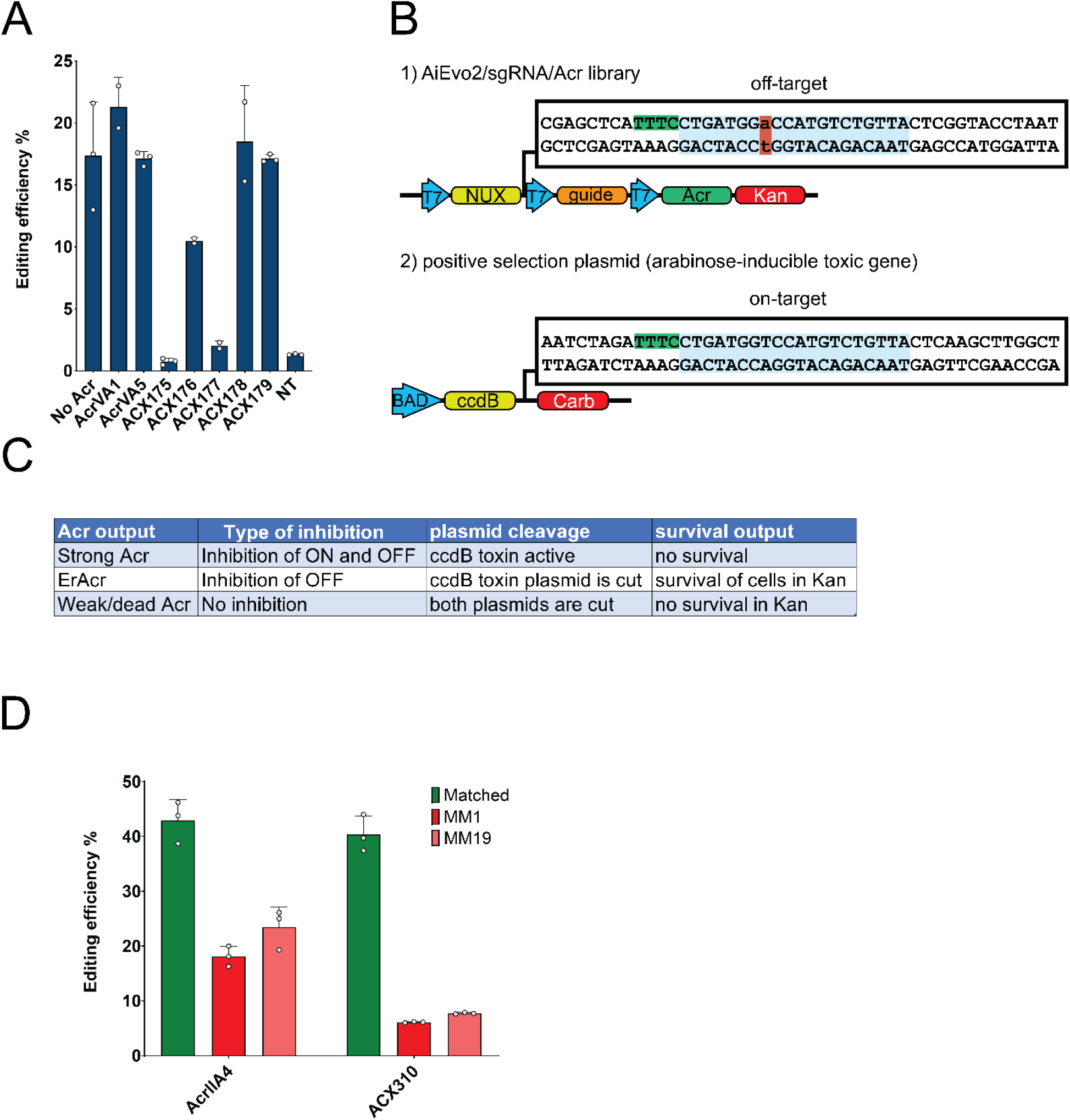
Discovery and engineering of precision enhancing ErAcr proteins for AiEvo2. **(A)** Distant homologues of AcrVA1 show strong inhibition of AiCas12a. Editing efficiencies are measured with sanger/TIDE and AiEvo2 targeting DNMT1. NT represents cells that were not transfected and were used as a no-editing control. **(B**) Schematic of ErAcr bacterial screening system. Plasmid 1 contains an off-target sequence and encodes AiEvo2, the ErAcr library, and a guide targeting DNMT1. Plasmid 2 contains the on-target site and encodes the ccdB toxin under arabinose induction. **(C)** Expected outcomes of the screen of ErAcrs. ErAcr variants will enable on-target cleavage of the ccdB toxin plasmid but inhibit off-target cleavage of Plasmid 1, allowing the bacteria to survive on both Kanamycin (Kan) and arabinose plates. **(D)** Editing efficiency of AiEvo2 in HEK293T cells targeting DNMT1. Fully matched on-target editing (green) is unaffected by ErAcr-310 relative to AcrIIA4 which does not inhibit Cas12a nucleases (p = 0.4). DNMT1 guides with mismatches at positions 1 (red) or 19 (pink) are significantly inhibited by ErAcr (p < 0.002). Editing was performed using plasmid transfection of HEK293T cells and analyzed by Sanger sequencing and TIDE analysis. N = 3 biological replicates.

With these strong AiEvo2 inhibitors in hand, we next sought to create engineered anti-CRISPR (ErAcr) proteins that would allow for on-target editing while fully inhibiting off-target editing. To do this, we modified a bacterial selection system previously used to screen for Cas mutants with increased specificity^23,24^. To ensure that the ErAcr variants inhibit off-target editing, we added an off-target site with a single mismatch as position 8 to the nuclease, guide, and ErAcr library expression plasmid (Figure 5B). Maintenance of on-target activity is selected for by the presence of a fully matched target site on a *ccdB* toxin plasmid under arabinose induction (Figure 5B). ErAcr variants that inhibit off-target editing but allow for on-target activity will survive on both Kanamycin and arabinose selection plates (Figure 5C). Several ErAcr variants that survived the screening conditions were selected for testing in human cells. The ErAcr variant ErAcr-310 was able to selectively inhibit mismatched off-target editing by AiEvo2 while maintaining on-target editing at the fully matched on-target site (Figure 5D).

## Discussion

The gene editing therapeutics field is experiencing the first wave of CRISPR therapies advancing through the clinic^35,54^. Along with this advancement, is the growing understanding of the importance of editing specificity. This has been highlighted by the numerous clinical holds on investigational new CRISPR therapies and the issuance of industry guidance by the FDA around detection and quantification of unintended off-target editing. The ability to precisely edit at nearly any genomic location while eliminating the risks of off-target editing outcomes is essential for new treatments for increasingly complex genetic conditions.

In this work, we identify and characterize AiCas12a, a novel type V CRISPR effector from *Acinetobacter indicus,* and demonstrate its activity in human cells. Native AiCas12a shares similar characteristics to other members of the Cas12a family, including a preference for TTTV PAM sequences, a PAM-distal staggered 5′ overhang cleavage site, and preference for a 20 nucleotide spacer. A key differentiator between AiCas12a and type II CRISPR effectors like SpyCas9 is the short sgRNA (40 nucleotides versus >100 for SpyCas9^55^) and the ability for AiCas12a to process a multiplexed guide array into individual guides capable of simultaneous editing at multiple genomic locations.

We optimized the NLS of AiCas12a for efficient human gene editing and further enhanced AiCas12a by engineering the nuclease to dramatically increase the PAM recognition, increase activity, and maintain the high intrinsic specificity of the Cas12a family. The resulting engineered enzyme, AiEvo2, is highly active, precise, and capable of targeting a high density of sites in the human genome. We show that AiEvo2 is able to selectively edit a single *HTT* allele in Huntington’s disease patient fibroblasts by discriminating a single base change between the two alleles. We then demonstrate highly efficient editing in primary hepatocytes after delivery of AiEvo2 mRNA encapsulated in LNP. Finally, we show that AiEvo2 can engineer effective CAR-T therapies by triplex editing and simultaneous CAR insertion in primary human T cells.

To further increase the precision of AiEvo2, we identified and engineered an anti-CRISPR protein to create an ErAcr capable of selectively inhibiting off-target editing while allowing for effective on-target therapeutic editing. We demonstrate efficient reduction of off-target editing in human cells using engineered ErAcr-310 with no reduction in on-target editing.

Together, these results demonstrate the efficacy and utility of AiEvo2 as a robust editor for a variety of human therapeutic applications. The ability to pair AiEvo2 with ErAcr-310 gives drug developers an additional tool for improving the safety profile of CRISPR therapies while maintaining the requisite efficacy of these systems. AiEvo2 and ErAcr-310 represent much needed additions to the CRISPR toolbox and are an important step toward safer gene editing therapies.

## Methods

### Nuclease discovery

The Cas12a candidate library was generated by searching known Cas12a sequences against the NCBI nonredundant (nr) and metagenomic (env_nr) protein databases using Basic Local Alignment Search Tool (BLAST)^56^. Additional candidates were identified through an hmmsearch^57^ of EBI protein databases using the Cpf1 hidden Markov model (HMM) from ^58^ as input. The genomic source sequences of the nucleases were then searched for CRISPR arrays and other Cas genes. CRISPR arrays were identified with CRISPRDetect ^59^ and a local version of CRISPRCasTyper ^60^. Known Cas genes were searched for using HMM profiles built from Pfam ^61^ seed sequences with the HMMer suite ^57^.

### Plasmid generation

All plasmids used for HEK293T tests of AiCas12a and Acr proteins were made by Twist Biosciences by synthesizing the nuclease CDS and cloning into a CMV expression vector from Twist Biosciences. Guide plasmids were cloned into a mammalian expression cassette bearing a U6 promoter upstream of the direct repeat followed by a BbsI cloning cassette and a termination poly-T sequence of TTTTTTgTTTT. Direct repest (DR) sequences were cloned downstream of the U6 promoter by PCR with oligos having the direct repeat sequences as overhangs and spacers were added based on the oligo anneal method using the BbsI restriction site. AiCas12a direct repeat sequences are listed in the Supplemental Data.

The plasmids used for the bacterial assay were modelled off previous reports^23,24,62,63^. Gblocks were ordered from Twist Biosciences and assembled using Gibson Assembly.

The plasmids for multiplex assays were made according to protocol described by ^19^ using oligo annealing PCR amplification.

### Plasmid transfections

Plasmids were transfected into HEK293T cells using Mirus Transit X2 reagent. 50,000 cells were plated per well of a 96 well plate and immediately transfected with 100 ng of nuclease expression vector and 50 ng of the corresponding sgRNA vector. When anti-CRISPR proteins were present, 100 ng of nuclease plasmid, 50 ng of sgRNA vector plasmid and 100 ng of Acr plasmid were transfected. Samples were incubated for 72 h and harvested with Quick Extract. Genomic DNA was amplified using KAPA HiFi polymerase with primers specific to the targeted genomic region. Amplicons were submitted for Sanger sequencing or next-generation sequencing (NGS). Editing was determined by analysis of ab1 files (Sanger) using TIDE batch ^64^ to calculate the editing efficiency of each sample. NGS files were analyzed using CRISPResso2^65^.

### PAM preference determination

PAM preferences were determined using the HD-PAMDA assay^11^. Briefly, a spacer capable of targeting a randomized PAM plasmid library made with 10-bp of randomized PAMs incorporated downstream of the SDR of the gRNA. The effective PAMs for the nuclease are depleted during the process, and the remaining PAMs are determined by NGS. Preferred PAM sequences for AiCas12a, AiEvo1, and AiEvo2 are listed in the Supplemental Data. Values are calculated based on ^11^ and PAM preferences are listed in order of preference.

### Cleavage site analysis

HEK293T cells were transfected with plasmids bearing AiCas12a and sgRNA targeting the FANCF locus in a 6 well dish following the Mirus transIT X2 (MIR6000) recommendations. Cells were incubated for 48h and harvested following an established protocol^11^. Each well from the plate was lysed in 100 uL of final volume. Lysates were aliquoted in 12uL portions and 10uL were used in a 20uL cleaving reaction. For the cleaving reaction, 10x cleaving buffer (5 mL of 1 M HEPES (pH 7.5), 15 mL of 5 M NaCl, 2.5 mL of 1 M MgCl2, 27.5 mL deionized water) 10uL of 1M DTT per mL. For a 20uL cleaving reaction, we add 50 ng of FANCF amplicon DNA (PCR amplified and cleaned) in 8uL and 2uL of 10x cleaving buffer with DTT. 10uL of the lysate aliquot is added into the mixture of 10uL of FANCF amplicon and cleaving buffer. Sample is incubated at 37C for 1 hour. After 1h, the reaction is quenched by 5uL of loading dye with SDS and incubated at 70C for 10 minutes. All the 20uL volume is loaded into a 2% agarose gel. Bands are extracted from the gel, cleaned and sequenced through Sanger with forward and reverse primers. Ab1 files are used to align the edges to the genomic sequence.

### HD patient cell editing

AiEvo2 protein was expressed in E. coli and purified using Ni-NTA chromatography (Genscript). AiEvo2 protein was complexed with guide RNA (IDT) for 20 minutes at 25°C. The resulting RNP complex was added to HD patient cells containing the rs363009 SNV (Coriell CM04723) and nucleofected using a Lonza 4D nucleofector system. Cells were allowed to incubate for 72 hours post-nucleofection before harvesting and genomic DNA extraction using Quick Extract solution. Editing efficiencies were determined using NGS as described above.

### LNP editing in primary mouse hepatocytes

mRNA encoding AiEvo2 (Northern RNA) and sgRNA targeting either the murine TTR or PCSK9 gene were encapsulated into GenVoy-ILM LNP (Precision Nanosystems) at 1:1 mRNA:sgRNA (w/w) and 4.5 N/P ratio. C57BL/6 mouse primary hepatocytes (Cell Biologics) were thawed and plated in 96 well tissue culture plates. The cells were incubated with LNP at 2.25ug of RNA per well. Post LNP transfection, the cells were allowed to incubate for 24 hours before harvesting and genomic DNA extraction. Editing efficiencies were determined using Sanger sequencing and TIDE analysis as described above.

### Primary T Cell editing and CAR-T generation

Human peripheral blood Pan-T cells (Stemcell Technologies) were thawed and expanded following manufacturer’s instructions. After 48 hours, the cells were transduced with adeno-associated virus serotype 6 (AAV6) containing a CD19 CAR integration cassette. Cells were then nucleofected with a mixture of three RNPs comprised of AiEvo2 and sgRNAs targeting TCRA, B2M, and PD-1. After 72 hours, cells were harvested for analysis. Genomic DNA was analyzed for editing by extracting edited CAR-T cells using Quick Extract solution. Editing efficiencies were determined using Sanger sequencing and TIDE analysis as described previously.

### Flow Cytometry analysis

1x10^6^ cells for each nucleofection condition were harvested and stained with a cocktail containing antibodies against TCRA, B2M and PD-1 as well as CD25 to check for activation. All antibodies used are pre-conjugated with fluorophores (BD biosciences: Hu CD3 FITC UCHT1 25Tst, Hu CD8 APC HIT8A 25Tst, Hu CD25 APC BC96 25Tst, Hu CD279 (PD-1) PE-Cy7 EH12.1 100Tst and Hu Bta2-Microglobulin PE TU99 100Tst). Cells were diluted to a concentration of 1-5x10^6^ cells/ml in ice cold FACS Buffer (PBS, 0.5-1% BSA) and stained in loBind 1.5mL tubes. 100uL of cell suspension were added to empty tubes on ice. 100 μl of Fc block was added to each sample (Fc block diluted in FACS buffer at 1:50 ratio). Samples were incubated on ice for 20 min and centrifuged at 1500 rpm for 5 min at 4°C. Supernatant was discarded and the recommended manufacturer’s concentration for the antibody cocktail was added. Samples were incubated in the dark at room temperature for 30 minutes. Cells were washed 3 times by centrifugation at 1500 rpm for 5 minutes and resuspended in 200 μl to 1ml of ice cold FACS buffer. 100 μl 1-4% paraformaldehyde was added to each sample and tubes were incubated for 10-15 min at room temperature. After the incubation, samples were spun at 1500 rpm for 5 min, fixing agent was removed, and cells were resuspended in 200 μl to 1 ml of ice cold PBS. Samples were analyzed on MACSquant analyzer 10.

### CAR insertion analysis

CD19 CAR insertion was assessed by extraction of mRNA from CAR-T cells, generation of cDNA, and quantitative analysis using qPCR. Fold expression of the CD19 CAR mRNA was calculated using the 2^-ΔΔCT method. Integration of the CD19 CAR was assessed through genomic DNA amplification using PCR primers targeting the inserted CAR cassette. The presence of a strong PCR product indicates integration of the CAR at the TCRA site.

### NALM6 cytotoxicity

CD19+ NALM6 cancer cells (NALM6, clone G5,CRL-3273**^™^**from ATCC) were incubated with CAR-T cells for 48 hours at effector to target (E:T) ratios of 0.1 to 2. Viable NALM6 cells were assessed by staining for CD-19 using antibodies followed by flow cytometry analysis as described above.

### ErAcr bacterial screen

The ErAcr bacterial screen was modeled off previous reports ^23,66,67^. E. coli strain BW25141 (DE3) were a kind gift from Dr. Edgel and were transformed with a plasmid containing a ccdB toxin induced by Arabinose as well as an on-target site for DNMT1. Single colonies were picked while in the presence of Carbenicillin. These were then replated in the presence of Arabinose to validate the correct toxicity phenotype. Electrocompetent cells were generated by washing log phase cells with 4 volumes of ice cold water and concentrating 100 fold in 10% glycerol. A mutant Acr library was made using ACX175 as a backbone and error-prone PCR with low mutation rate using GeneMorph II random mutagenesis kit (Agilent, Santa Clara, CA, USA). The resulting library was subcloned into a second plasmid containing AiEvo2, the sgRNA targeting the DNMT1 sequence and an off-target target DNMT1 sequence. Surviving colonies were selected and sequenced. Candidate ErAcrs were transferred to a mammalian expression vector downstream of a CMV promoter and were tested in HEK293T cells with AiEvo2 and sgRNA targeting DNMT1 as well as DNMT1 with mismatches on position 1 and 19.

## Supporting Information

Supporting Information: Figures depicting guide structures, editing activity, CAR-T schematics and flow cytometry, and protein alignment.

## Acknowledgements

We would like to thank Prof. Joseph Bondy-Denomy for critical reading of the article.

