## Supplemental Information for "Discovery and engineering of AiEvo2, a novel Cas12a nuclease for human gene editing applications"

### Supporting Information

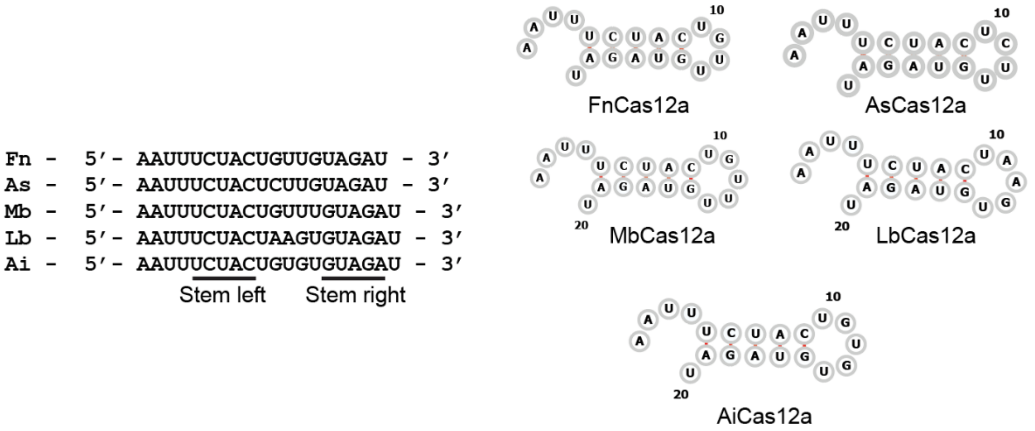


**Figure S1.** Direct repeat sequences of commonly used Cas12a nucleases and AiCas12a showing the predicted RNA structures on RNAFold [10]. MbCas12a, LbCas12a and AiCas12a form a 7-nucleotide loop as opposed to a 6-nucleotide loop formed by AsCas12a and FnCas12a direct repeats.

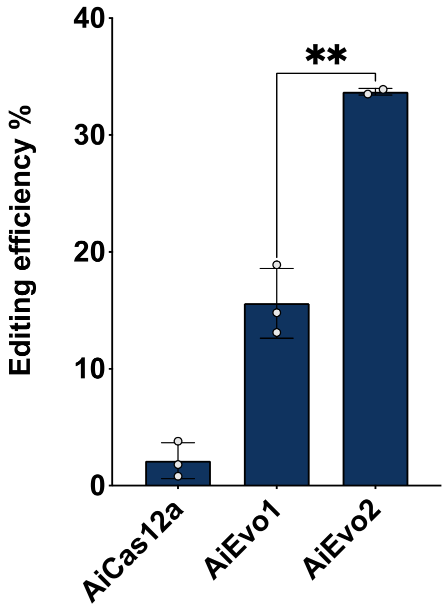


**Figure S2.** Comparison of wild-type AiCas12a, AiEvo1, and AiEvo2 targeting the DNMT1 locus. Editing was performed using plasmid transfection in HEK293T cells and analyzed by Sanger sequencing and TIDE analysis ($p = 0.004$). N=3 biological replicates.

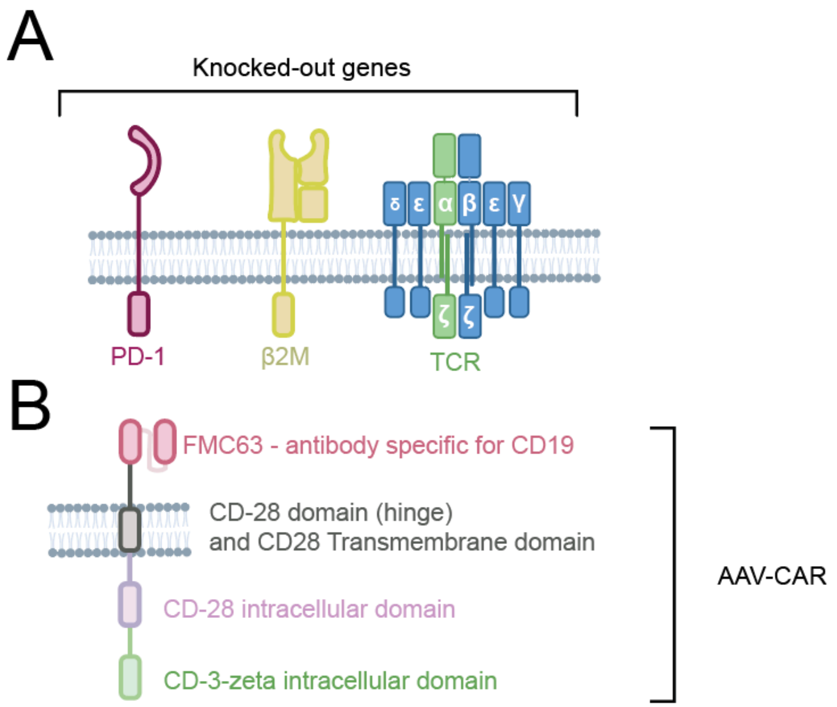


**Figure S3. (A)** Schematic of the CAR-T relevant targets that were knocked out by AiEvo2 and **(B)** final AAV based CAR design

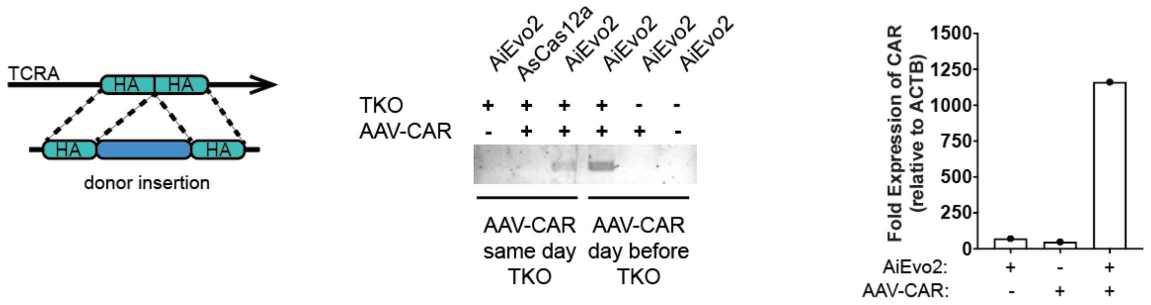


**Figure S4.** Confirmation of CAR integration. Schematic and PCR on genomic DNA showing integration of the CAR in the TCRA locus. CAR expression relative to ACTB is confirmed by qPCR using the ΔΔCT method.

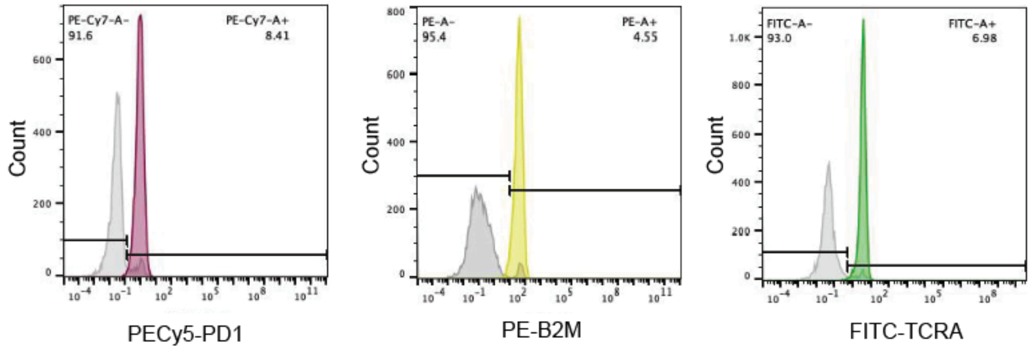


**Figure S5.** Flow cytometry measurement of triple-knockout T cells showing the loss of expression of each surface marker for the deleted genes. All targets had less than 9% of residual expression.

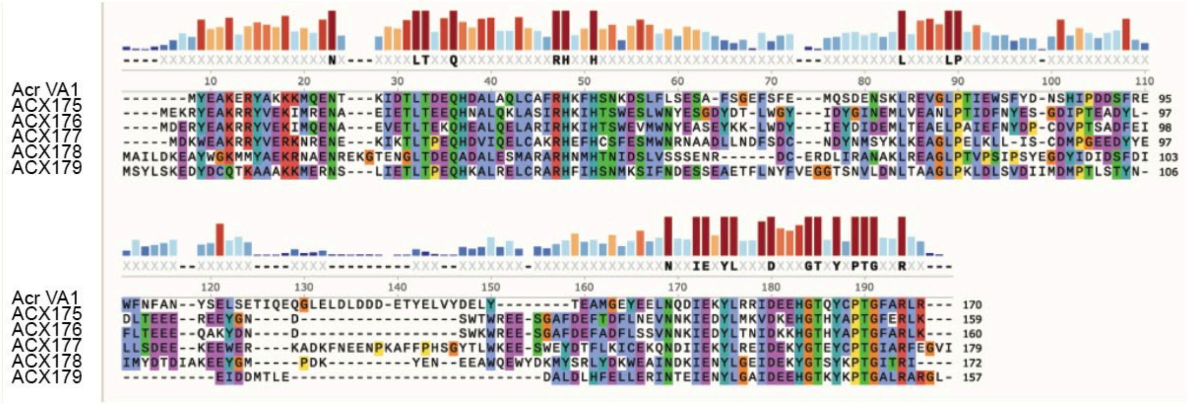


**Figure S6.** Amino acid alignment of AcrVA1 and Cas12a Acr candidates.
